# The combined treatment of Molnupiravir and Favipiravir results in a marked potentiation of antiviral efficacy in a SARS-CoV-2 hamster infection model

**DOI:** 10.1101/2020.12.10.419242

**Authors:** Rana Abdelnabi, Caroline S. Foo, Suzanne J. F. Kaptein, Xin Zhang, Lana Langendries, Laura Vangeel, Judith Breuer, Juanita Pang, Rachel Williams, Valentijn Vergote, Elisabeth Heylen, Pieter Leyssen, Kai Dallmeier, Lotte Coelmont, Dirk Jochmans, Arnab K. Chatterjee, Steven De Jonghe, Birgit Weynand, Johan Neyts

## Abstract

Favipiravir and Molnupiravir, orally available antivirals, have been reported to exert antiviral activity against SARS-CoV2. In recent days preliminary efficacy data have been reported in COVID-19 patients. We here studied the combined antiviral effect of the drugs in the SARS-CoV2 hamster infection model. We first demonstrate that Molnupiravir can reduce infectious virus titers in lungs of infected animals in a dose-dependent manner by up to 3.5 log_10_ which is associated with a marked improvement of virus-induced lung pathology. When animals are treated with a combination of suboptimal doses of Molnupiravir and Favipiravir (that each alone result in respectively a 1.3 log_10_ and 1.1 log_10_ reduction of infectious virus titers in the lungs), a marked combined potency is observed. Infectious virus titers in the lungs of animals treated with the combo are on average reduced by 4.5 log_10_ and infectious virus are no longer detected in the lungs of 60% of treated infected animals. Both drugs result in an increased mutation frequency of the remaining viral RNA recovered from the lungs. In the combo-treated hamsters an increased frequency of C-to-T and G-to-A mutations in the viral RNA is observed as compared to the single treatment groups which may explain the pronounced antiviral potency of the combination. Our findings may lay the basis for the design of clinical studies to test the efficacy of the combination of Molnupiravir and Favipiravir in the treatment of COVID-19.

## Main text

The severe acute respiratory syndrome coronavirus 2 (SARS-CoV-2) was first identified in Wuhan, China in December 2019 ^1^. Since then, the virus rapidly spread around the globe with more than 115 million cases and 2.5 million deaths reported until 3^rd^ March 2021 [www.covid19.who.int]. Infection with SARS-CoV-2 results in coronavirus-induced disease (COVID-19) which is characterized by a wide range of symptoms including fever, dry cough, muscle and/or joint pain, headache, decreased sense of taste and smell and diarrhea. The disease can progress into severe complications such as acute respiratory distress syndrome (ARDS), respiratory failure, septic shock as well as multi-organ failure, which are mainly attributed to a massive cytokine storm and exaggerated immune response ^2^.

To date, there are no approved, selective coronavirus antivirals to treat or prevent infections. Even those vaccinated may not all be protected against infection and disease, in particular following infection with variants that are less susceptible to the current vaccines. Antivirals against SARS-CoV-2 will, at least when given early enough after a positive test or after onset of symptoms, reduce the chance to progress to (more) severe disease. In addition, such antiviral drugs will be useful to protect for example healthcare workers and high-risk patients in a prophylactic setting. Such drugs are also needed for the treatment of immunodeficient patients who do not mount a (sufficiently robust) immune response following vaccination. Since the *de novo* development and approval of (a) specific, highly potent antiviral(s) for SARS-CoV-2 may require years, the main focus for COVID-19 treatment in the current pandemic is to repurpose drugs that have been approved or in clinical trials for other diseases ^3^.

We recently demonstrated that the anti-influenza drug Favipiravir results in a pronounced antiviral activity in SARS-CoV-2-infected hamsters ^4^. The efficacy of the drug is currently being explored in multiple phase II clinical studies. Interim data from a multicenter, open-labeled study of Avifavir (Favipiravir) in patients with COVID-19, reveals a faster virological response, a shorter time to clinical symptoms and reduced mortality rates compared to patients receiving the Standard of Care (Corritori et al., Science Spotlight™, Conference on Retroviruses and Opportunistic Infections [CROI] 2021). The ribonucleoside analogue, N4-hydroxycytidine (NHC, EIDD-1931), was initially developed as an influenza inhibitor, but exerts also broader-spectrum antiviral activity against multiple viruses belonging to different families of RNA viruses. Activity against SARS-CoV-2 has been reported in cell lines and primary human airway epithelial cell cultures ^5^.

Both Favipiravir and NHC act through lethal mutagenesis. Incorporation into viral RNA results in the accumulation of deleterious transition mutations beyond a permissible error threshold to sustain the virus population, leading to error catastrophe ^6,7^. The orally bioavailable, pro-drug counterpart of NHC ^8^, Molnupiravir (EIDD-2801, MK-4482) was shown to result in an antiviral effect against SARS-CoV-2 in a Syrian hamster model ^9^, in a mouse model ^10^ and in ferrets ^11^. Data from a first-in-human, phase 1, randomized, double-blind, placebo-controlled study in healthy volunteers indicate that the drug is well tolerated and that plasma exposures exceed the expected efficacious doses based on scaling from animal models ^12^. The drug is currently being assessed for its potential as an antiviral treatment of SARS-CoV-2 infection in Phase 2 clinical trials of infected patients (NCT04405570, NCT04405739). Very recently interim data (on one secondary objective) were reported in 202 non-hospitalized adults who had signs or symptoms of COVID-19 and active confirmed SARS-CoV2 infection. At day 5, a reduction in positive viral culture from nasopharyngeal swabs was noted in subjects who received molnupiravir (0/47) as compared to placebo (6/25). No safety signals were identified in the 202 participants (Painter et al., Science Spotlight™, CROI 2021).

We here report on the pronounced combined activity of Favipiravir and Molnupiravir in the hamster infection model.

## Results

### Dose-response efficacy of Molnupiravir against SARS-CoV-2 in Syrian hamsters

We first evaluated the (dose-response) effect of Molnupiravir (EIDD-2801) in SARS-CoV-2-infected hamsters to select an appropriate dose for the combination study. Briefly, 6-8 weeks female SG hamsters were treated orally with Molnupiravir (either 75, 150, or 200 mg/kg, BID) or the vehicle (i.e. the control group) for four consecutive days starting one hour before intranasal infection with SARS-CoV-2. At day four post-infection (pi), the animals were euthanized and lungs were collected for quantification of viral RNA, infectious virus titers and lung histopathology as described before^4^ (Fig. 1A). Molnupiravir treatment resulted in a dose-dependent reduction in the viral RNA copies per mg of lung tissue with 1.3 (P=0.002), 1.9 (P<0.0001) and a 3.3 log_10_ (P<0.0001) reduction was noted in the groups that had been treated BID with 75, 150 or 200 mg/kg, respectively (Fig. 1B). A similar pattern was observed for the infectious virus load in the lungs whereas the intermediate and high doses, but not the 75 mg/kg dose BID, significantly reduced infectious virus lung titers (Fig. 1C). The reduction in infectious virus titers (TCID_50_/mg tissue) in the lungs of hamsters treated BID with 150 and 200 mg/kg was 1.3 (P=0.0002) and 3.5 (P<0.0001) log_10_, respectively (Fig. 1C). Treatment with 75, 150 and 200 mg/kg Molnupiravir BID significantly reduced the histological lung disease score (P=0.0025, P=0.005, P<0.0001, respectively) (Fig. 1D). All the doses studied were well tolerated without significant weight loss or any obvious adverse effects (Fig. 1E).

**Fig. 1.**
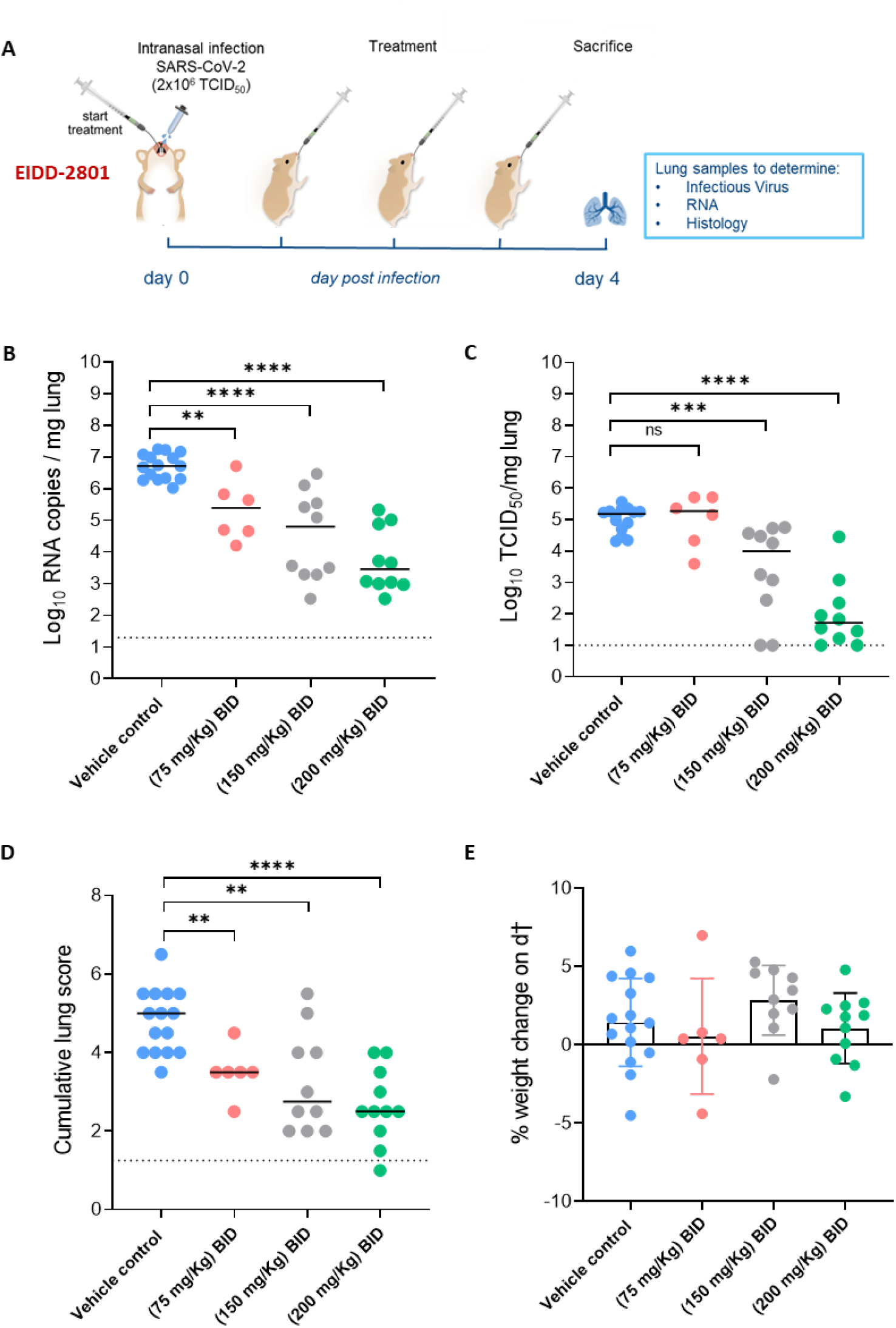
Dose-response efficacy of Molnupiravir (EIDD-2801) against SARS-CoV-2 in a hamster infection model. (A) Set-up of the study. (B) Viral RNA levels in the lungs of control (vehicle-treated) and EIDD-2801-treated (75, 150 or 200 mg/kg, BID) SARS-CoV-2-infected hamsters at day 4 post-infection (pi) are expressed as log_10_ SARS-CoV-2 RNA copies per mg lung tissue. Individual data and median values are presented. (C) Infectious viral loads in the lungs of control (vehicle-treated) and EIDD-2801-treated SARS-CoV-2-infected hamsters at day 4 pi are expressed as log_10_ TCID_50_ per mg lung tissue Individual data and median values are presented. (D) Cumulative severity score from H&E stained slides of lungs from control (vehicle-treated) and EIDD-2801-treated SARS-CoV-2-infected hamsters. Individual data and median values are presented and the dotted line represents the median score of untreated non-infected hamsters. (E) Weight change at day 4 pi in percentage, normalized to the body weight at the time of infection. Bars represent means ± SD. Data were analyzed with the Mann-Whitney U test. *P < 0.05, **P < 0.01, ***P < 0.001, ****P < 0.0001, ns=non-significant. All data (panels B, C, D, E) are from two independent experiments except for the 75 mg/kg group. The number of animals were 15, 6, 10, and 10 for respectively the vehicle 75, 150, and 200 mg/kg condition.

### The combined treatment of Molnupiravir and Favipiravir results in a marked potentiation of efficacy

Next, we studied what the combined efficacy is of suboptimal doses of Molnupiravir and Favipiravir (Fig. 2A). Treatment with Favipiravir alone (300 mg/kg, BID) reduced viral RNA and infectious virus loads in the lungs of infected animals by 0.7 (P=0.0009) and 1.2 (P=0.0002) log_10_/mg tissue, respectively (Fig. 2B/C). Treatment with Molnupiravir (150 mg/kg BID) only resulted in 1.9 (P<0.0001) and 1.3 (P=0.0002) log_10_/mg reduction in viral RNA and infectious virus loads respectively. The combined treatment resulted in a reduction of 2.7 log_10_ viral of lung viral RNA titers (Fig. 2B), but interestingly, in a markedly enhanced reduction in infectious virus titers (4.5 log_10_ TCID_50_ per mg lung, P=0.02, P=0.0005 as compared to Molnupiravir and Favipiravir alone, respectively) (Fig. 2C). Notably, there was no detectable infectious virus in the lungs of six out of ten hamsters in the combined treatment group (Fig. 2C). A marked improvement in the histological lung pathology scores was also observed in the combined treatment group (Fig. 2D) and no significant weight loss or toxicity signs were observed (Supplementary Fig. S1).

**Fig. 2.**
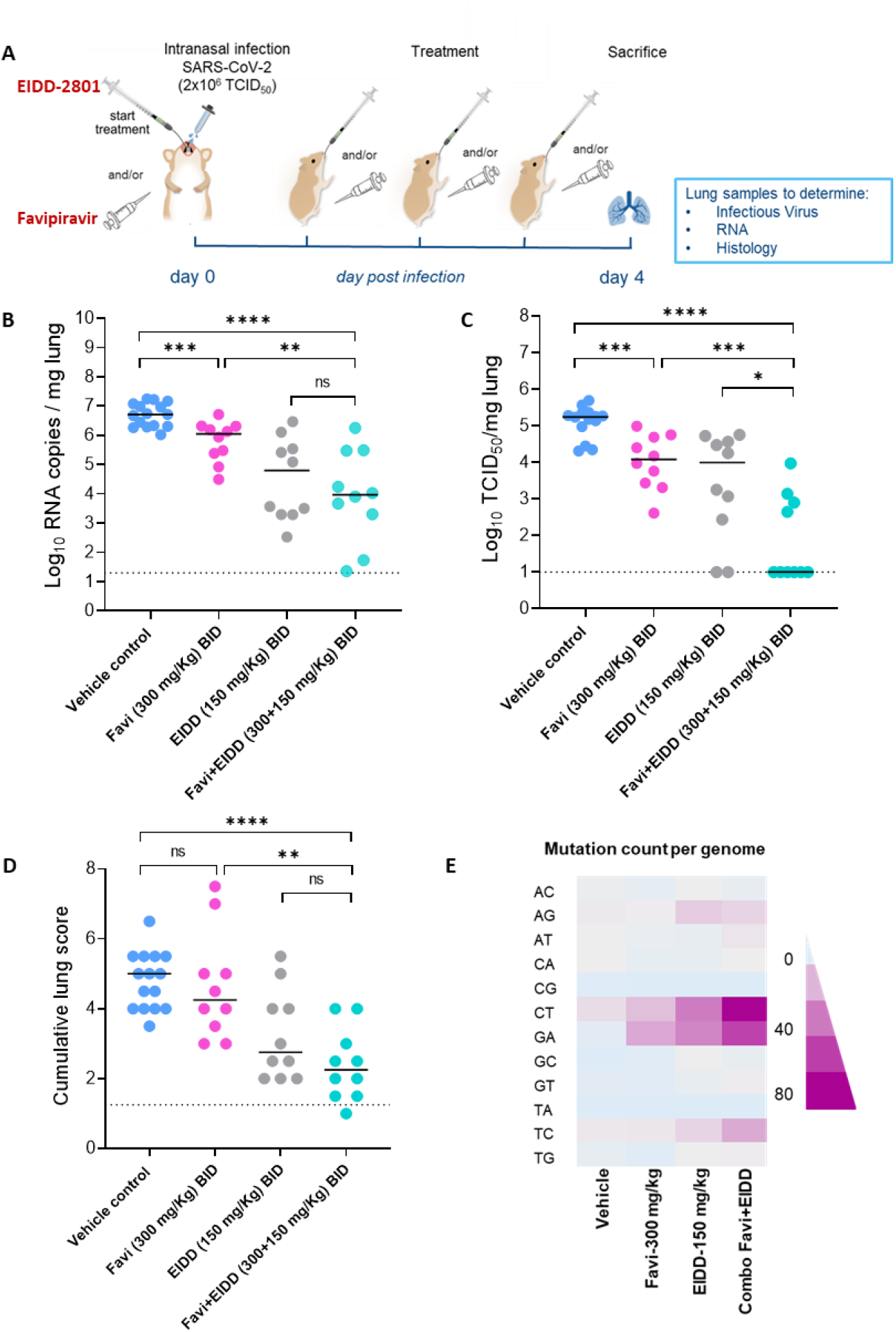
Combined efficacy of Favipiravir and Molnupiravir (EIDD-2801) against SARS-CoV-2 in a hamster infection model. (A) Set-up of the study. (B) Viral RNA levels in the lungs of control (vehicle-treated), Favipiravir-treated (300 mg/kg, BID), EIDD-2801-treated (150 mg/kg, BID) and combination-treated (Favipiravir+EIDD-2801 at 300+150 mg/kg, BID, respectively) SARS-CoV-2-infected hamsters at day 4 post-infection (pi) are expressed as log_10_ SARS-CoV-2 RNA copies per mg lung tissue. Individual data and median values are presented. (C) Infectious viral loads in the lungs of control (vehicle-treated), Favipiravir-treated, EIDD-2801-treated and combination-treated (Favipiravir+EIDD-2801) SARS-CoV-2-infected hamsters at day 4 pi are expressed as log_10_ TCID_50_ per mg lung tissue. Individual data and median values are presented. (D) Cumulative severity score from H&E stained slides of lungs from control (vehicle-treated), Favipiravir-treated, EIDD-2801-treated and combination-treated (Favipiravir+EIDD-2801) SARS-CoV-2-infected hamsters. Individual data and median values are presented and the dotted line represents the median score of untreated non-infected hamsters. (E) Mean mutation count (per the whole genome) in the viral RNA isolated from the lungs of control (vehicle-treated), Favipiravir-treated (300 mg/kg, BID), EIDD-2801-treated (150 mg/kg, BID) and combination-treated (Favipiravir+EIDD-2801 at 300+150 mg/kg, BID, respectively) SARS-CoV-2-infected hamsters at day 4 post-infection (pi). Data were analyzed with the Mann-Whitney U test. *P < 0.05, **P < 0.01, ***P < 0.001, ****P < 0.0001, ns=non-significant. Favi=Favipiravir, EIDD=EIDD-2801. All data (panels B, C, D) are from two independent experiments with 15, 10, 10 and 10 animals for respectively the vehicle, Favipiravir 300 mg/kg, EIDD-2801 150 mg/kg and Favipiravir+EIDD-2801 condition.

Molnupiravir is known to increase the mutation frequency of MERS-CoV viral RNA in infected mice ^5^. To test whether this is also the case in SARS-CoV-2-infected hamsters, we used Illumina deep sequencing to determine the SARS-CoV-2 mutations rate in remaining viral RNA in lung samples of hamsters after treatment. A dose-dependent increase in the mutation count (in particular, C-to-T and G-to-A transitions) in samples from Molnupiravir treated hamsters was observed as compared to the vehicle control group (Supplementary Fig. S2). The Molnupiravir (150mg/kg) + Favipiravir (300 mg/kg) combination resulted in a markedly higher number of C-to-T and G-to-A mutations (68 and 50, respectively), as compared to the single dose groups [150 mg/kg Molnupiravir group (33 and 31, respectively) and 300 mg/kg Favipiravir (14 and 21, respectively)] (Fig. 2E). The C-to-T and G-to-A mutation count in the combination group was also markedly higher than in the highest dose group of Molnupiravir (45 and 39, respectively) (Supplementary Fig. S2). These results may at least partially explain the markedly enhanced reduction in infectious viral loads observed in the combination treatment group.

## Discussion

Remdesivir (Veklury), is the first drug to have received FDA approval for use in hospitalised COVID19 patients, although the World Health Organisation has issued a conditional recommendation against its use on 20 Nov 2020 (www.who.int). Remdesivir needs to be administrated intravenously which precludes its use in the early stages of the infection/disease or even a prophylactic use. We previously demonstrated that treatment of SARS-CoV-2-infected hamsters with a high doses of Favipiravir largely reduces infectious virus titers in the animals and results as a consequence in a markedly improved lung pathology ^4^. Favipiravir is currently being studied for the treatment of COVID-19 in clinical trials in several countries and was very recently reported to result in COVID-19 patients in a faster virological response, a shorter time to clinical symptoms and reduced mortality rates (Corritori et al., Science SpotlightTM, CROI 2021). Also Molnupiravir has been reported to exert therapeutic and prophylactic activity against SARS-CoV-2 in several animal models ^9–11^. Importantly very recent interim data from phase II clinical studies in COVID-19 patients, revealed a reduction in the time required to reach negative isolation of infectious virus from the nasopharyngeal swabs from participants with symptomatic SARS-CoV-2 infection (Painter et al., Science Spotlight™, CROI 2021). Both Molnupiravir and Favipiravir have a high barrier to resistance, resistant variants have a loss in fitness and induce lethal viral mutagenesis ^6,7^.

We here demonstrate that the combination of suboptimal doses of Molnupiravir and Favipiravir results in a marked antiviral activity in the hamster model. Infectious virus titers were reduced to undetectable levels in 6 out of 10 treated animals. A median reduction of 4.5 log_10_ TCID_50_/mg lung tissue was achieved, which is markedly more pronounced than what could be expected from an additive activity of either Molnupiravir (1.3 log_10_) or Favipiravir (1.1 log_10_) when dosed alone. This pronounced efficacy of the combination may partially or even entirely be explained by the increased total mutation count in viral RNA collected from the lungs of combo-treated hamsters as compared to the single treatment groups.

In conclusion, the combination of Molnupiravir and Favipiravir (two oral drugs with a high barrier to resistance for which there is very recent initial evidence that they exert antiviral activity in COVID-19 patients), is particularly effective in the treatment of SARS-CoV2 infections in hamsters. This efficacy may be explained by an enhanced accumulation of mutations as compared to monotherapy. Our findings may lay the basis for the design of clinical studies to test the efficacy of the combination of Molnupiravir and Favipiravir in the treatment of COVID-19.

## Methods

### SARS-CoV-2

The SARS-CoV-2 strain used in this study, BetaCov/Belgium/GHB-03021/2020 (EPI ISL 109 407976|2020-02-03), was recovered from a nasopharyngeal swab taken from an RT-qPCR confirmed asymptomatic patient who returned from Wuhan, China in the beginning of February 2020. A close relation with the prototypic Wuhan-Hu-1 2019-nCoV (GenBank accession 112 number MN908947.3) strain was confirmed by phylogenetic analysis. Infectious virus was isolated by serial passaging on HuH7 and Vero E6 cells ^4^; passage 6 virus was used for the study described here. Live virus-related work was conducted in the high-containment A3 and BSL3+ facilities of the KU Leuven Rega Institute (3CAPS) under licenses AMV 30112018 SBB 219 2018 0892 and AMV 23102017 SBB 219 20170589 according to institutional guidelines.

### Cells

Vero E6 cells (African green monkey kidney, ATCC CRL-1586) were cultured in minimal essential medium (Gibco) supplemented with 10% fetal bovine serum (Integro), 1% L-glutamine (Gibco) and 1% bicarbonate (Gibco). End-point titrations were performed with medium containing 2% fetal bovine serum instead of 10%.

### Compounds

For the first pilot experiment, EIDD-2801 was kindly provided by Calibr at Scripps Research (USA). For further studies, Molnupiravir (EIDD-2801) was purchased from Excenen Pharmatech Co., Ltd (China) and was formulated as 50 or 100 mg/ml (for groups with the highest dose) stocks in a vehicle containing 10%PEG400 and 2.5% Kolliphor-EL in water. Favipiravir was purchased from BOC Sciences (USA) and was formulated as a 50 mg/mL stock in 3% sodium bicarbonate.

### SARS-CoV-2 infection model in hamsters

The hamster infection model of SARS-CoV-2 has been described before ^4,13^. Housing conditions and experimental procedures were approved by the ethics committee of animal experimentation of KU Leuven (license P065-2020). For infection, female hamsters of 6-8 weeks old were anesthetized with ketamine/xylazine/atropine and inoculated intranasally with 50 μL containing 2×10^6^ TCID_50_ SARS-CoV-2 (day 0).

### Treatment regimen

For dose-response treatment, animals were treated twice daily with 75, 150 or 200 mg/kg of EIDD-2801 by oral gavage just before infection with SARS-CoV-2. For combination therapy, hamsters were treated from day0 with 150 mg/kg EIDD-2801 (oral gavage) and 300 mg/kg Favipiravir (intraperitoneal, i.p.) twice daily. All the treatments continued until day 3 pi. Hamsters were monitored for appearance, behavior and weight. At day 4 pi, hamsters were euthanized by i.p. injection of 500 μL Dolethal (200mg/mL sodium pentobarbital, Vétoquinol SA). Lungs were collected and viral RNA and infectious virus were quantified by RT-qPCR and end-point virus titration, respectively.

### SARS-CoV-2 RT-qPCR

Hamster lung tissues were collected after sacrifice and were homogenized using bead disruption (Precellys) in 350 μL TRK lysis buffer (E.Z.N.A.^®^ Total RNA Kit, Omega Bio-tek) and centrifuged (10.000 rpm, 5 min) to pellet the cell debris. RNA was extracted according to the manufacturer’s instructions. RT-qPCR was performed on a LightCycler96 platform (Roche) as described before ^4^.

### End-point virus titrations

Lung tissues were homogenized using bead disruption (Precellys) in 350 μL minimal essential medium and centrifuged (10,000 rpm, 5min, 4°C) to pellet the cell debris. To quantify infectious SARS-CoV-2 particles, endpoint titrations were performed as described before ^4^.

### Histology

For histological examination, the lungs were fixed overnight in 4% formaldehyde and embedded in paraffin. Tissue sections (5 μm) were analyzed after staining with hematoxylin and eosin and scored blindly for lung damage by an expert pathologist. The scored parameters, to which a cumulative score of 1 to 3 was attributed, were the following: congestion, intra-alveolar hemorrhagic, apoptotic bodies in bronchus wall, necrotizing bronchiolitis, perivascular edema, bronchopneumonia, perivascular inflammation, peribronchial inflammation and vasculitis.

### Deep sequencing and analysis of whole genome sequences

Genomic sequences from all samples were obtained using SureSelect^XT^ target enrichment and Illumina sequencing. Reads generated were trimmed with Trim Galore (https://github.com/FelixKrueger/TrimGalore). Duplicated reads were removed using Picard (http://broadinstitute.github.io/picard). Reads from the inoculation sample were mapped to the SARS-CoV-2 reference genome (NC_045512) from GenBank using BWA-MEM ^14^. The mapping quality was checked using Qualimap and the consensus whole genome sequence was generated using QUASR ^15,16^. Reads from the lung samples were mapped to this unique reference sequence. Genomes with less than less than a 100 read depth were excluded. Variants above 1% and with a minimum of 2 supporting reads per strand were identified at sites with a read depth of ≥ 10 using VarScan ^17^.

### Statistics

GraphPad Prism (GraphPad Software, Inc.) was used to perform statistical analysis. Statistical significance was determined using the non-parametric Mann Whitney U-test. P-values of ≤0.05 were considered significant.

## Data Availability

All of the data generated or analysed during this study are included in this published article.

## Acknowledgments

We thank Carolien De Keyzer, Lindsey Bervoets, Thibault Francken, Elke Maas, Jasper Rymenants, Birgit Voeten, Dagmar Buyst, Niels Cremers, Bo Corbeels and Kathleen Van den Eynde for excellent technical assistance. We are grateful to Piet Maes for kindly providing the SARS-CoV-2 strain used in this study. We thank Prof. Jef Arnout and Dr. Annelies Sterckx (KU Leuven Faculty of Medicine, Biomedical Sciences Group Management) and Animalia and Biosafety Departments of KU Leuven for facilitating the animal studies. This project has received funding from the Covid-19-Fund KU Leuven/UZ Leuven and the COVID-19 call of FWO (G0G4820N), the European Union’s Horizon 2020 research and innovation program under grant agreements No 101003627 (SCORE project) and Bill & Melinda Gates Foundation (BGMF) under grant agreement INV-00636. Sequencing was performed by the Pathogen Genomics Unit at UCL. R.A., C.S.F. and L.L. were supported by a KU Leuven internal project fund. X.Z. received funding of the China Scholarship Council (grant No.201906170033). J.B. receives funding from the NIHR UCL/UCLH Biomedical Research Centre. J.P. is funded by the Rosetrees Trust.

## Author Contributions

R.A., C.S.F. and J.N. designed the studies; R.A., C.S.F., S.J.F.K., X.Z., J.B., J.P. and L.L. performed the studies; R.A., C.S.F., J.B, J.P. and B.W. analyzed data; D.J. and J.N. provided advice on the interpretation of data; R.A., C.S.F. and J.N. wrote the paper with input from co-authors; A.K.C. and S.D.J provided essential reagents; V.G. and E.H. provided and facilitated access to essential infrastructure; R.A., C.S.F., R.W. and J.N. supervised the study; L.V., L.C., P.L., J.N. and K.D. acquired funding.

---

**Fig. S1.**
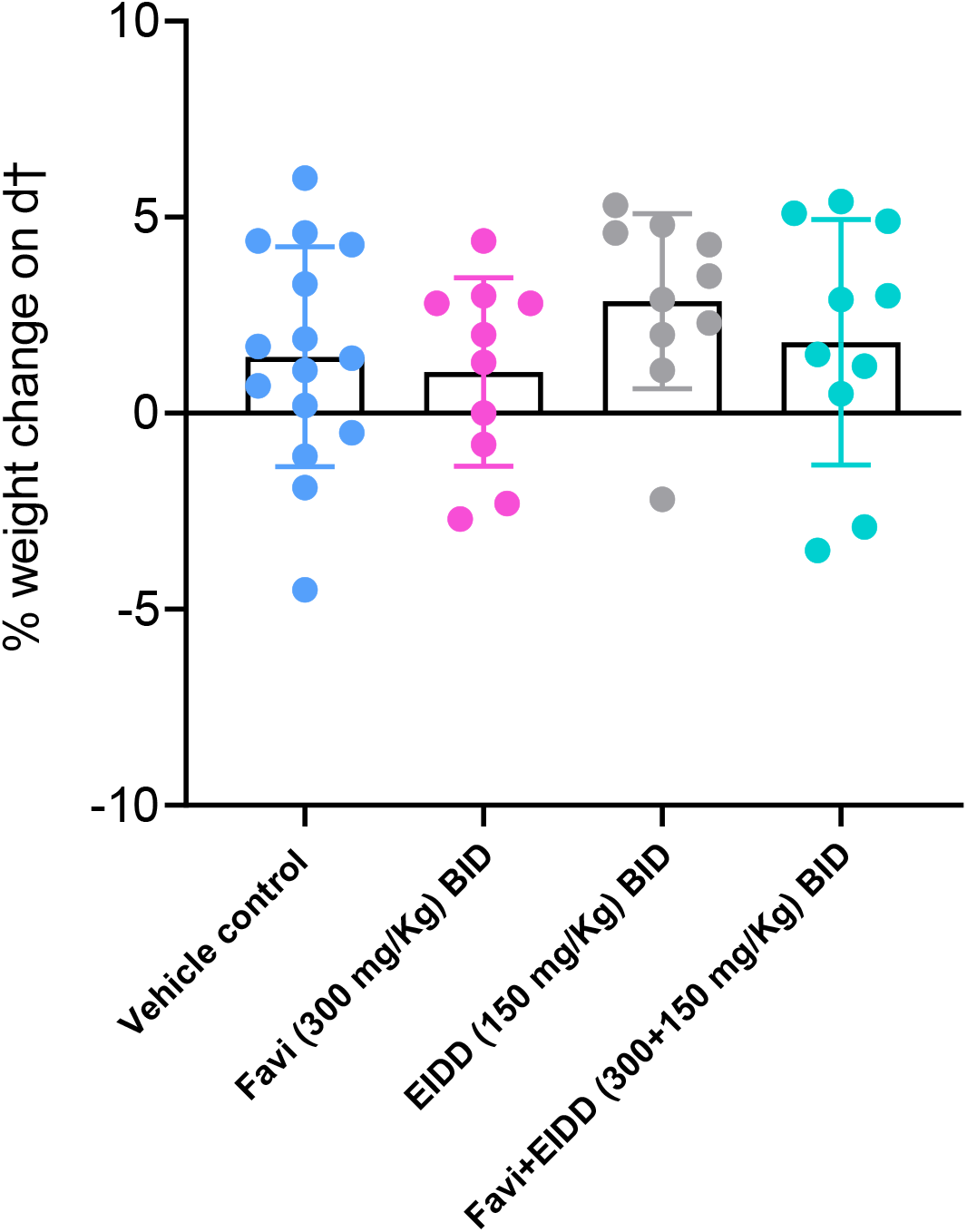
Tolerability of combined treatment with Favipiravir and EIDD-2801 in SARS-CoV-2-infected hamsters. Weight change at day 4 post-infection in percentage, normalized to the body weight at the time of infection. Bars represent means ± SD.

**Fig. S2.**
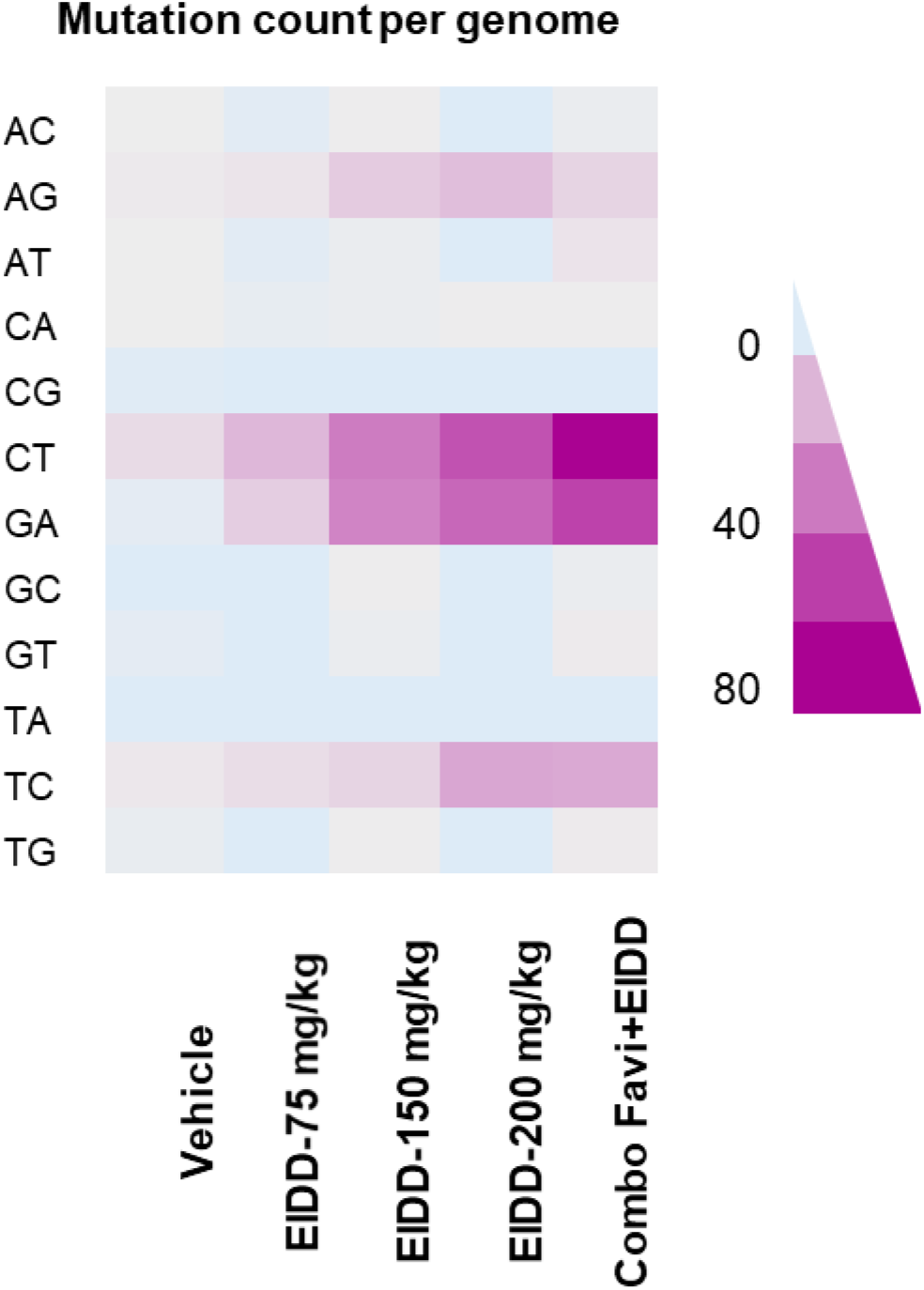
Dose-dependent increase in mutation count in SARS-CoV-2 viral RNA by Molnupiravir (EIDD-2801). Mean mutation count (per the whole genome) in the viral RNA isolated from the lungs of control (vehicle-treated), EIDD-2801-treated (75, 150 or 200 mg/kg, BID) and combination-treated (Favipiravir+EIDD-2801 at 300+150 mg/kg, BID, respectively) SARS-CoV-2-infected hamsters at day 4 post-infection (pi).

